# Ripples precede insight in the right human hippocampus

**DOI:** 10.64898/2026.02.05.703313

**Authors:** Kotaro Yamashiro, Takamitsu Iwata, Yuji Ikegaya, Yasushi Iimura, Takumi Mitsuhashi, Hiroharu Suzuki, Katsuyuki Negishi, Takuya Suematsu, Ryosuke Yamamoto, Ryohei Fukuma, Naoki Tani, Hui Ming Khoo, Satoru Oshino, Haruhiko Kishima, Takufumi Yanagisawa

## Abstract

Insight is a sudden transition from uncertainty to awareness of a solution, often preceded by non-conscious processing. Despite the ubiquity of insights in human cognition, the neural mechanisms underlying this pre-conscious restructuring phase remain largely unknown. Recent theoretical frameworks suggest that the hippocampus, through its capacity for rapid associative retrieval and flexible recombination of stored representations, may facilitate the non-conscious exploration that precedes insight. Hippocampal ripples associated with memory replay and consolidation, represent a promising candidate mechanism for this process. We recorded intracranial local field potentials from depth electrodes in the hippocampus of ten patients with drug-resistant epilepsy while they performed non-verbal reasoning tasks. Participants indicated the moment of solution awareness via button press, enabling precise temporal alignment of neural activity to subjective insight. We observed a significant increase in ripple rate during a pre-response window. Critically, this effect was lateralized to the right hippocampus. These findings provide the first direct human electrophysiological evidence linking right-lateralized hippocampal ripples to the non-conscious phase of insight. We propose that right hippocampal ripples support rapid retrieval and flexible recombination of representations, potentially triggering cortical processes that culminate in conscious solution awareness.

## Introduction

Insight is marked by a sudden transition from uncertainty to conscious solution awareness, frequently accompanied by abruptness and high confidence (Knoblich et al. 1999; Kounios and Beeman 2014). Psychological theories of insight propose that sudden solutions arise from a change in how a problem is mentally represented. After reaching an impasse, individuals overcome unhelpful assumptions by restructuring the problem, allowing previously hidden relationships to become apparent (Kaplan and Simon 1990; Knoblich et al. 1999). This restructuring can involve constraint relaxation and recombination of prior knowledge such that the solution is experienced as sudden even if preparatory processes unfold outside consciousness (Knoblich et al. 1999; Kounios and Beeman 2014). In the present study, we define insight as the abrupt emergence of conscious solution awareness.

Neuroscientific work suggests that insight emerges from coordinated dynamics within a distributed network spanning temporal and prefrontal cortices, with right-hemisphere contributions frequently implicated in semantic integration and recognition of distantly related associations (Bowden and Jung-Beeman 2003; Jung-Beeman et al. 2004; Kounios and Beeman 2014). Using association tasks that can yield insight versus non-insight solutions, functional magnetic resonance imaging (fMRI) and electroencephalogram (EEG) studies report distinct neural signatures for insight solutions, including changes that arise close to and sometimes preceding the subjective moment of solution awareness (Jung-Beeman et al. 2004; Sheth, Sandkühler, and Bhattacharya 2009). Causal evidence also points to a role of the right anterior temporal lobe in facilitating insight on compound association problems (Salvi et al. 2020). Collectively, these findings support the notion that insight is preceded by identifiable pre-solution neural processes that culminate in a sudden conscious breakthrough.

Evidence suggests that subcortical structures, particularly the hippocampus may also contribute to the representational restructuring that precedes insight by enabling rapid binding and recombination of relational elements. Consistent with this view, hippocampal engagement has been reported in human neuroimaging studies of insight-like comprehension and problem solving (Aziz-Zadeh, Kaplan, and Iacoboni 2009; Luo and Niki 2003). These results motivate the hypothesis that hippocampal computations may help generate candidate relational structures that, once sufficiently coherent, can enter awareness as an insight solution.

Hippocampal ripples, brief high-frequency population events, provide a candidate physiological mechanism linking hippocampal computations to insight. In rodents, ripples support hippocampo–cortical communication and temporally compressed reactivation and recombination of neuronal sequences that contribute to memory consolidation and planning (Buzsáki 2015). In humans, ripples have been linked to internally oriented cognition and network dynamics (Kaplan and Simon 1990), and to memory processes such as encoding- and recall-related activity (Norman et al. 2019, 2021). Recent work further connects ripples to naturally occurring self-generated thought, supporting the idea that ripples accompany internal simulation and integrative cognition beyond externally cued memory tasks (Iwata et al. 2024). Despite these advances, it remains unknown whether ripples contribute to the neural processes that precede and enable the moment of insight during active problem-solving.

In the present study, we tested whether hippocampal ripples contribute to pre-solution processes leading to insight in humans. We recorded intracranial local field potentials (LFPs) from hippocampal depth electrodes in patients with drug-resistant epilepsy while they performed non-verbal reasoning tasks from the Wechsler Adult Intelligence Scale (WAIS). Participants indicated solution awareness via button press, providing a precise temporal marker for insight onset as defined above. We hypothesized that hippocampal ripple rates would increase in a pre-solution window immediately preceding the button press, consistent with the idea that ripple-associated hippocampal computations support rapid retrieval and recombination of relational elements that enable the sudden emergence of a coherent solution into awareness.

## Materials and Methods

### Participants and intracranial recordings

The study cohort comprised patients with drug-resistant epilepsy who underwent intracranial electrode implantation for the presurgical evaluation of seizure onset zones and memory function. Depth electrodes were implanted in the hippocampus or parahippocampal gyrus based solely on clinical necessity to localize epileptogenic regions. To minimize potential selection bias, the EEG data analyst was not involved in the determination of electrode placement. Recordings were excluded from analysis if the hippocampus was pathologically diagnosed with hippocampal sclerosis to minimize the influence of underlying pathology on the neurophysiological data.

### Study Population and Ethics

Of the thirteen participants consented to study procedures, data from three participants were excluded since interictal epileptiform discharges (IEDs) were observed during the trial, leaving data from ten participants for the final analysis. The sample consisted of 8 males and 2 females, with 7 right-handed, 1 left-handed, and 2 participants with unreported handedness. Among the participants, 4 patients had electrodes implanted bilaterally, 4 had electrodes implanted in the left hemisphere, and 2 had electrodes implanted in the right hemisphere. Data collection was conducted at the Epilepsy Center of Juntendo University Hospital. The research protocol was approved by the Institutional Review Board (approval no.18-164), and written informed consent was obtained from all participants prior to their inclusion in the study.

### Task design

To characterize the electrophysiological correlates of insight in non-verbal reasoning, we employed two subtests from the WAIS: Matrix Reasoning and Picture Completion (Hartman 2009; Lichtenberger and Kaufman 2009). The Matrix Reasoning task, a measure of fluid reasoning, presented participants with incomplete visual pattern matrices and required the selection of a missing piece to complete the logical sequence. The Picture Completion task, assessing visual perception and attention to detail, required participants to identify a specific missing feature within an image of an object or scene. The experimental protocol for each task consisted of two practice trials followed by 25 test problems presented in order of increasing difficulty. Crucially, to time-lock neural activity to the moment of subjective insight, participants were instructed to press a button immediately upon realizing the solution (Figure 1A). Following the button press, they indicated their answer; if an incorrect response was provided, participants were permitted to re-attempt the problem until the correct solution was identified.

**Figure 1.**
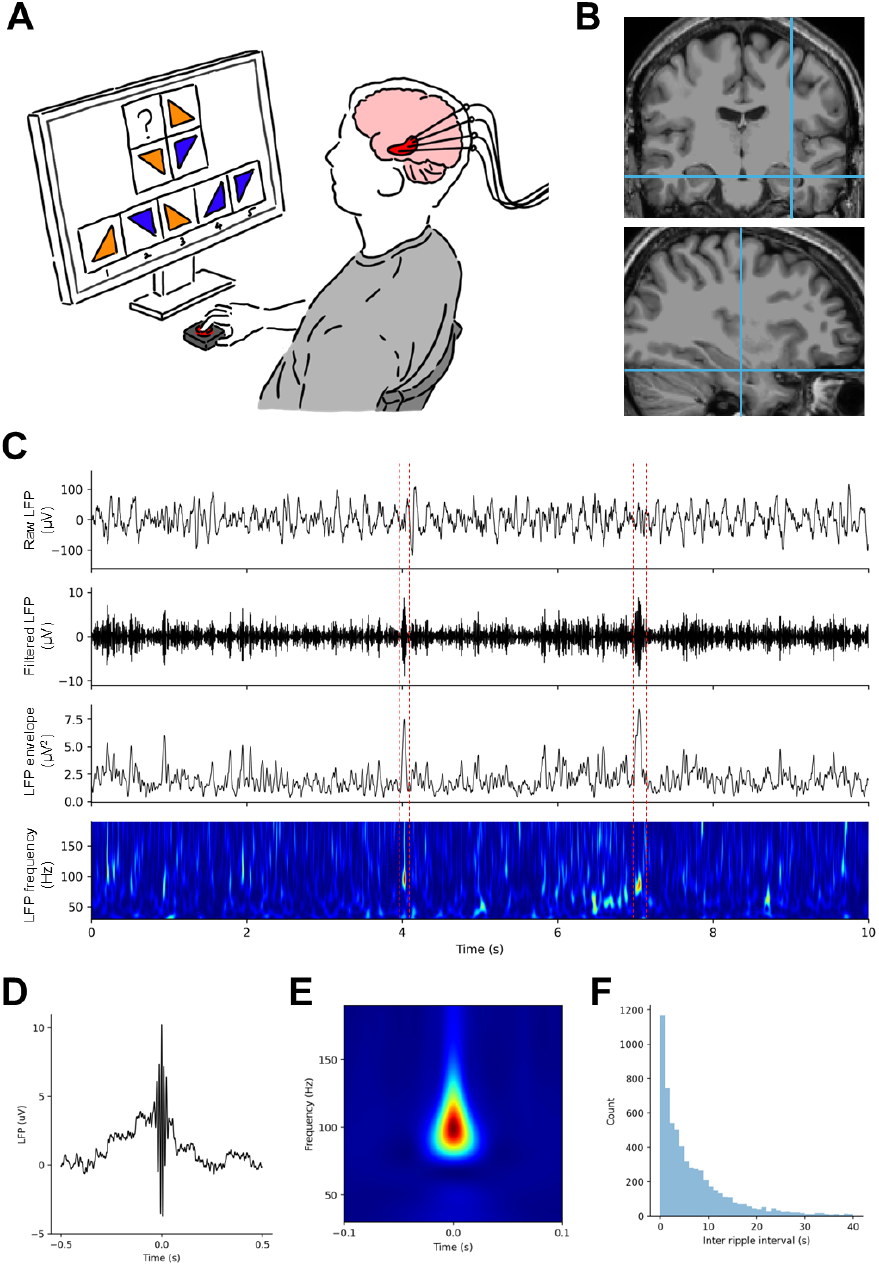
Experimental Setup and Analysis of LFP Signals during Tasks. (A) Schematic of the experimental setup. The participant was seated in front of a monitor while intracranial EEG signals were recorded during task performance. The tasks included Matrix Reasoning and Picture Completion. (B) Sample MRI image with coordinate of the electrode in the hippocampus. (C) Example LFP trace showing: (i) the raw bipolar signals, (ii) the filtered LFP signals in the 30–190 Hz band, (iii) the signal envelope, and (iv) the wavelet transform. (D) Global average of ripples across all trials. (E) Global average of the wavelet transform applied to all ripples. (F) Histogram of inter-ripple intervals across trials.

### Data Acquisition

All data were acquired in the hospital rooms during the participants’ inpatient stay at Juntendo University Hospital. To mitigate the acute effects of surgery, only EEG data recorded from 4 days post-implantation onward were included in the analysis. Data collection was conducted in a controlled environment, ensuring that external noise or disturbances did not interfere with the recordings.

During the problem-solving task, vocal responses were captured using a microphone attached to the participant’s collar, which was synchronized with the EEG acquisition system to ensure precise temporal alignment of verbal outputs. The problem-solving task required participants to press the button with their dominant hand when they thought they knew the answer, with accuracy and response time recorded as behavioral measures of task performance.

Intracranial EEG data were recorded from depth electrodes using an EEG-1200 system (Nihon Kohden, Tokyo, Japan) with a sampling rate of 10 kHz. Signal preprocessing employed a low-pass filter with a cutoff frequency of 3000 Hz and a high-pass filter at 0.016 Hz. The depth electrodes featured an intercontact spacing ranging from 5 to 15 mm and a contact width of 1 mm, strategically placed to cover key areas of the hippocampus and other regions of interest (Figure 1B).

Electrode localization was performed by coregistering postoperative computed tomography (CT) scans with preoperative T1-weighted magnetic resonance imaging (MRI) images using SYNAPSE VINCENT software (Fujifilm, Tokyo, Japan). MRI data were acquired using a SIGNA Architect (GE Healthcare) and CT imaging was obtained using a Discovery CT750 HD (GE Healthcare).

### Data preprocessing

Prior to ripple analysis, data were downsampled to 2 kHz using an 8th-order Chebyshev Type I low-pass filter with a cutoff frequency of 900 Hz. Prior to ripple analysis, all recorded EEG data were visually inspected, and periods containing seizures or intense epileptic activity were identified and discarded. IEDs were detected using established methods (Smith et al. 2022). Briefly, raw iEEG signals were band-pass filtered between 80 and 140 Hz with a zero-phase (two-pass), fourth-order Butterworth filter. Events exceeding eight times the standard deviation of the full recording session were labeled as IEDs, and recordings were additionally reviewed to identify subthreshold IEDs. Of the 13 participants who consented to participate, 3 exhibited IEDs during the session and were excluded from subsequent analyses. Channels with persistent muscle artifact, line noise, or other technical disturbances were also removed. After quality control, 54 hippocampal channels were retained, including 20 from the right hemisphere and 34 from the left hemisphere.

### Ripple detection

Hippocampal ripples were detected following a set of criteria adapted from previous literature (Iwata et al. 2024; Kunz et al. 2024). Ripples were detected from bipolar signals from hippocampal contacts. For participants who were implanted bilaterally, hippocampal channels from both hemispheres were included in the analysis. The bipolar signals were band-pass filtered between 80 and 140 Hz using a two-pass, 4th-order Butterworth filter. The instantaneous analytic amplitude of the signal was then computed using the Hilbert transform, and these amplitudes were smoothed with a 20 ms smoothing window to reduce noise and improve detection accuracy. The mean and standard deviation of the smoothed amplitude were calculated across the entire recording. Candidate ripple events were defined as time periods where the signal exceeded 2 standard deviations above the mean amplitude (Figure 1C).

Candidate ripple events were retained as physiological ripples only if they met all the following criteria: (1) the peak smoothed amplitude exceeded 3 standard deviations above the mean, (2) the event duration was longer than 20 ms and shorter than 500 ms, (3) the band-pass filtered signal contained at least three peaks and three troughs, and (4) the normalized power spectrum exhibited a global peak between 80–140 Hz. The power spectrum was computed from 30–190 Hz in 2 Hz steps using Morlet wavelets with seven cycles. It was then divided by the power spectrum estimated across the entire recording, ensuring that the ripple events had a characteristic spectral signature.

### Ripple features and phase locking to delta wave

To characterize the detected ripple events, we quantified several basic features for each ripple and summarized their distributions across all events. Specifically, we measured (i) event duration (time from ripple onset to offset), (ii) ripple amplitude, defined as the peak of the smoothed analytic amplitude of the 80–140 Hz band-pass–filtered signal, (iii) peak ripple frequency, estimated as the frequency at which spectral power was maximal within the ripple band during the event, and the number of oscillatory cycles contained within each ripple event.

To examine the relationship between hippocampal ripples and ongoing delta activity, we computed the delta phase associated with each ripple. Bipolar hippocampal signals were band-pass filtered in the delta range (0.5–2 Hz) using an 8,000-order Butterworth filter. The instantaneous phase of the delta-band signal was then estimated using a Hilbert transform. For each ripple, we extracted the delta phase at the ripple peak time (*i*.*e*., the time of maximum smoothed ripple-band analytic amplitude). We then assessed the overall phase relationship between ripples and delta oscillations by averaging ripple-locked delta phases across events (and, where applicable, across channels/participants) to quantify whether ripples preferentially occurred at specific phases of the delta rhythm. To test whether ripple occurrence exhibited statistically significant phase locking to the delta oscillation, we applied a Rayleigh test and a second-harmonic Rayleigh test.

### Peri-response ripple analysis

Behavioral responses were first classified as correct or incorrect based on whether the subsequent answer matched the WAIS solution key. The button press indicating solution awareness was treated as the response onset (time 0) and used to align ripple events across trials. Peri-response ripple rates were computed separately for the Matrix Reasoning and Picture Completion tasks, and further stratified by hemisphere (left vs. right hippocampus) and accuracy (correct vs. incorrect). For each hippocampal bipolar channel, ripple times were expressed relative to the button press, and ripple rate was computed in non-overlapping 500-ms bins spanning −4 s to +4 s around the response. Within each bin, ripple rate was defined as the number of ripples occurring in the bin divided by bin duration (0.5 s), yielding an event rate (Hz). For visualization and summary statistics, ripple-rate time courses were averaged within each condition, with variability quantified as the standard error of the mean.

To assess whether any peri-response modulation exceeded what would be expected by chance alignment to the response, we generated surrogate ripple-rate traces for each condition by disrupting the temporal relationship between ripple times and response onset while preserving the overall temporal structure of the ripple train. This surrogate procedure was repeated 1,000 times to generate a null distribution. For each surrogate realization, peri-response ripple rates were recomputed using the identical binning procedure (500-ms bins; −4 to +4 s). Statistical significance of peri-response rate modulation was assessed by comparing the observed ripple rate in each time bin against the corresponding surrogate distribution. Resulting p-values across time bins were corrected for multiple comparisons using false discovery rate (FDR) correction.

## Results

### Hippocampal ripples during task

To characterize hippocampal ripple activity during task performance, we analyzed LFPs recorded from bipolar signals localized to the hippocampus in ten patients with drug-resistant epilepsy. Across participants, we detected 8,372 hippocampal ripple events (Figure 1D–F). We first quantified basic ripple features across all detected events (Figure 2A–D). Ripples had a mean duration of 36.35 ± 23.63 ms (mean ± standard deviation), a mean peak-to-peak amplitude of 44.44 ± 119.55 μV, a mean peak frequency of 87.49 ± 6.19 Hz, and contained 3.00 ± 2.25 oscillatory cycles. These distributions were consistent with the expected short-duration, narrowband, high-frequency oscillatory profile of hippocampal ripples.

**Figure 2.**
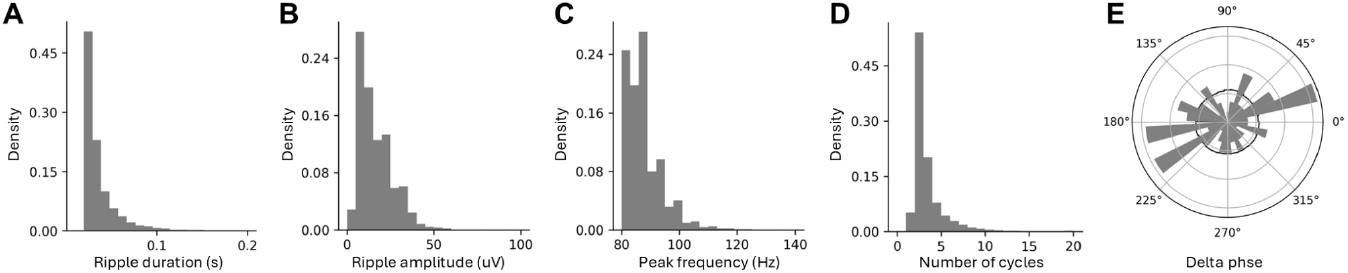
Characteristics of detected hippocampal ripples and their relationship to the delta phase. (A–D) Distributions across all detected ripples showing (A) event duration, (B) ripple amplitude (peak smoothed analytic amplitude of the 80–140 Hz band-pass–filtered signal), (C) peak ripple frequency, and (D) number of oscillatory cycles per event. (E) Ripple-locked delta phase (0.5–2 Hz) at the time of each ripple peak, shown for each hippocampal channel. Gray histograms show the empirical phase distribution, and the black trace shows the corresponding distribution from time-shuffled surrogate ripples (see Methods).

To examine whether ripples preferentially occurred at specific phases of the ongoing hippocampal delta rhythm, we extracted the instantaneous delta phase (0.5–2 Hz) at the time of each ripple peak (Figure 2E). Visual inspection suggested that ripples tended to occur near transitions between delta phases (i.e., around the rising/falling portions of the delta cycle), motivating formal tests for phase locking. We first assessed unimodal phase concentration using the circular mean and a standard Rayleigh test. The circular mean phase was 196.68°, but phase locking was not significant (Rayleigh test: *z* = 0.0525, *p* = 0.774), indicating no strong unimodal preference. Because ripple occurrence can plausibly cluster at two opposing phases of a slow oscillation, we next tested for antiphasic (bimodal) phase locking using a second-harmonic Rayleigh test. This analysis revealed significant biphasic clustering with a binormal circular mean at 36.63° and 216.63° (second-harmonic Rayleigh test: *z* = 0.239, *p* = 0.00480). Together, consistent with previous results, these results suggest that ripple timing was not locked to a single preferred delta phase, but instead showed antiphasic phase organization, consistent with ripple occurrence being modulated by hippocampal delta-cycle state (Weiss et al. 2020).

### Hippocampal ripple dynamics prior to insight

We next tested whether ripple occurrence was temporally modulated around the moment participants indicated solution awareness. Ripple events were aligned to the button press (time 0), and peri-response ripple rates were computed in 500-ms bins over a window spanning −4 to +4 s. Statistical significance was assessed by comparing the observed peri-response ripple-rate time course to a surrogate null distribution generated by 1,000 shuffles that disrupted the alignment between ripple times and response onset while preserving the temporal structure of the ripple train. P-values across time bins were corrected for multiple comparisons using FDR.

In the Matrix Reasoning task, ripple rates in the right hippocampus showed a significant peri-response increase on correct trials, exceeding the surrogate distribution from approximately −1.5 s to +0.5 s relative to the button press. No time bins survived correction on incorrect trials, although the peri-response trace showed a qualitatively similar increase. In contrast, left hippocampal electrodes did not show a significant peri-response increase relative to surrogates for either correct or incorrect trials (Figure 3A). A similar pattern was observed in the Picture Completion task. Ripple rates in the right hippocampus were significantly elevated relative to surrogates on correct trials from approximately −2.0 s to +0.5 s, whereas no bins reached significance for incorrect trials despite a comparable trend. Again, left hippocampal electrodes did not show significant peri-response modulation relative to surrogates for either accuracy condition (Figure 3B). Together, these results indicate that hippocampal ripple rates increase in the seconds preceding the behavioral report of solution awareness, with this effect selectively observed in the right hippocampus and most robust on correct trials.

**Figure 3.**
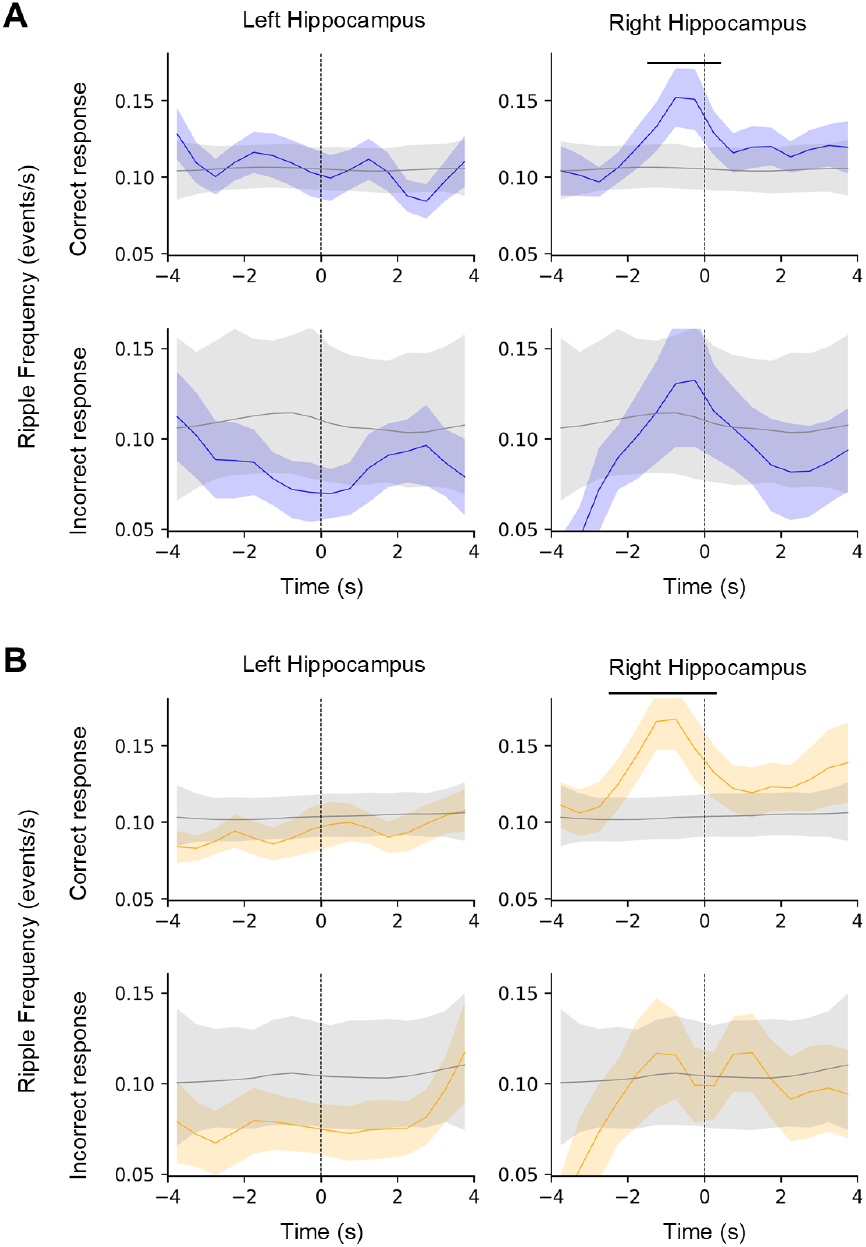
Ripple rate during response onsets. (**A**) Ripple rate computed for the Matrix Reasoning task in 500 ms time bins from -4s to +4s around the button press. Left panels show ripple rates from electrodes in the left hippocampus, while right panels show ripple rates from electrodes in the right hippocampus. Top panels display ripple rates for correct responses, and bottom panels show ripple rates for incorrect responses. The colored solid line represents the average trace with SEM indicated by the shaded area, while the gray solid line shows shuffled data with the 95% confidence interval in the shaded area. (**B**) Same as panel A, but for the Picture Completion task.

## Discussion

In this study, we provide the first direct human electrophysiological evidence that hippocampal ripples contribute to insight in problem-solving. By recording LFPs from depth electrodes in patients with drug-resistant epilepsy, we captured hippocampal activity with high temporal precision during non-verbal reasoning tasks. Participants indicated the moment of solution awareness via button press, providing a precise behavioral marker for the transition from uncertainty to conscious insight. We found a significant increase in ripple rate in the right hippocampus during a pre-solution window, specifically for correctly solved trials. Critically, this effect was absent in the left hippocampus, revealing a hemispheric asymmetry in the neural dynamics preceding insightful breakthrough. This work bridges a critical gap between cognitive theories of insight, which emphasize unconscious associative processing, and the neurophysiological mechanisms that enable such processing.

A critical methodological consideration in clinical intracranial recordings is the distinction between physiological ripples and pathological high-frequency activity, including IEDs and pathological ripples associated with epileptogenic tissue. Recent consensus guidelines emphasize that ripple detection in humans is highly sensitive to electrode localization, detection thresholds, and the specific criteria used to differentiate genuine ripples from other fast oscillations (Liu et al. 2022; Maslarova et al. 2025). We addressed these concerns through several complementary approaches. First, we excluded recording sessions containing detectable IEDs, ensuring that analyzed epochs were free from overt epileptiform activity. Second, we employed a stringent, multi-criterion detection algorithm that assessed not only band-limited amplitude (80–140 Hz) but also event duration, spectral peak characteristics, and oscillatory structure. Third, we examined the phase-locking relationship between detected ripples and low-frequency delta oscillations, a feature of physiological ripples in both rodent and human hippocampus (Weiss et al. 2020). The consistent phase-locking observed in our dataset provides strong evidence that the detected events represent physiological ripples rather than pathological high-frequency activity. Together, these methodological safeguards support the interpretation that the pre-solution ripple modulation reflects a genuine cognitive process rather than epilepsy-related artifact.

In rodents, ripples are highly synchronous hippocampal events that occur during offline states characterized by temporally compressed reactivation of neuronal ensembles, a phenomenon termed “replay” (Buzsáki 2015; Carr, Jadhav, and Frank 2011; Diba and Buzsáki 2007; Jadhav et al. 2012). Ripples can also express prospective sequences termed “preplay” that represent potential future trajectories or simulate new courses of action, suggesting a role in combining known and new information (Diba and Buzsáki 2007; Karlsson and Frank 2009; Pfeiffer and Foster 2013; Singer et al. 2013). We believe that the peri-solution ripple increase observed may reflect an analogous process adapted for abstract problem-solving: moments when hippocampal networks transiently engage replay-like computations to rapidly retrieve, recombine, and evaluate candidate conceptual associations (Joo and Frank 2018). The resulting restructuring would then cross the threshold into awareness, manifesting as the subjective experience of sudden insight (Kotler et al. 2025).

A striking feature of our findings is the hemispheric specificity of the pre-solution ripple modulation, which was evident in the right hippocampus. This lateralization is consistent with established evidence for functional specialization within the human medial temporal lobe (MTL), where left MTL systems preferentially engage with verbalizable, linguistic materials, whereas right MTL systems contribute more strongly to nonverbal, visuospatial, and relational representations (Chen et al. 2021; Golby 2001; Iglói et al. 2010). In spatial navigation, for instance, right hippocampal activity has been linked to allocentric place-based representations and configural processing (Iglói et al. 2010). The task employed in our study is a nonverbal reasoning task that relies on the detection of abstract relational structures. Given this task design, the preferential engagement of right hippocampal ripples is compatible with the computational demands of the problem-solving process. We believe that right hippocampal ripples may facilitate rapid retrieval and flexible binding of semantic associations that are distributed across right temporal and parietal cortical networks.

Several limitations constrain interpretation of our findings and suggest directions for future work. First, electrode placement was dictated by clinical considerations, resulting in heterogeneous coverage across participants. Although our between-subject results indicate right hippocampal specificity, broader bilateral coverage would allow more definitive characterization of hemispheric asymmetries. Second, solution awareness was indexed by button press, providing an objective temporal marker but likely introducing variable delays due to metacognitive judgment and motor execution. This temporal imprecision may blur neural dynamics preceding conscious awareness; finer alignment could be achieved using continuous behavioral measures or electrophysiological markers of decision formation. Future studies could address these limitations by incorporating eye-tracking and pupillometry to provide continuous indices of attention and arousal, which may also refine ripple detection given their association with low-arousal states (Liu et al. 2022). Beyond correlational analyses, combining balanced bilateral recordings with trial-by-trial phenomenological reports and causal perturbation approaches—such as closed-loop stimulation triggered by detected ripples—would enable stronger tests of whether hippocampal ripples play a causal role in insight. Finally, extending this paradigm to diverse insight tasks varying in modality, relational complexity, and knowledge demands would clarify the conditions under which hippocampal ripples support problem-solving and assess the generality of the observed right lateralization.

In conclusion, this study provides novel evidence of the role of hippocampal ripples, specifically in the right hippocampus, in guiding insight. The findings highlight the lateralized role of the hippocampus in integrating prior knowledge with incoming information to facilitate the transition from uncertainty to solution realization. These results advance our understanding of the neural mechanisms of insight and offer new evidence into how the hippocampus contributes not only to memory formation but also to insight.

## Acknowledgments

This work was supported by the Japan Science and Technology Agency (JST), Moonshot R&D-MILLENNIA Program (JPMJMS2012), ERATO (JPMJER1801), K program (JPMJKP25Y7), Japan Agency for Medical Research and Development (AMED) (JP24wm0625207), Sumino Isamu foundation.

We would like to express our deepest gratitude to Dr. György Buzsáki for his invaluable discussions and insights on hippocampal ripples in humans. We also wish to acknowledge the clinical staff at the Juntendo University Hospital for their exceptional support in patient care and assistance with the surgical procedures and electrode implantation. Additionally, we thank all the participants who generously contributed to this study. Without their participation, this work would not have been possible.

## Data availability

The data that support the findings of this study are available from the corresponding author upon reasonable request.

